# Croconium Dye Thermosensitive Liposomes for Light Activated Drug Release

**DOI:** 10.64898/2026.04.25.720841

**Authors:** James Kelley, Natalie Wehrle, Sarah Wessel, Yoonjee Park

**Affiliations:** University of Cincinnati

## Abstract

This study investigates a novel light-activated drug delivery system designed to produce on-demand drug release. The light-activated system was developed by incorporating a photostable photothermal agent, croconium dye, into liposomes to enable thermally triggered drug release. The drug release from the liposomes was determined at three powers of 210, 295, and 380 mW under 0-, 1-, and 2-minute light irradiation. A continuous wave 808 nm laser was used as the light source. Dexamethasone sodium phosphate (DSP) released from the liposomes was tunable depending on the power and irradiation time with a range of 1 -19 μg released depending on irradiation power and time. For local temperature measurement during the photothermal activation, polymerized 10, 12 – Pentacosadiynoic acid (PCDA) was incorporated in the lipid bilayer. Under heating polymerized PCDA undergoes a transition into a red phase from a blue phase. Utilizing the spectrum changes under known temperatures a regression model was developed to calculate the local temperature of the liposomes under irradiation. The ability of the liposomes to release DSP under irradiation in the presence of a phantom tissue was tested under different attenuation coefficients to match various common biological tissues. The liposomes were still able to release DSP in the presence of tissue phantoms for a certain thickness of the tissue. Finally, the cytotoxicity of the liposomes with the croconium dye for chemical and thermal toxicity was determined. The liposomes displayed good biocompatibility with Human Microvascular Endothelial Cell line-1 (HMEC-1). The results support the use of croconium dye as a potential alternative to commonly photothermal agents used in drug delivery such as metal nanoparticles. Future work will focus on optimization of absorbance spectrum for drug release, and in vivo studies for efficacy and safety.

## 1 INTRODUCTION

Liposomes are a common drug carrier for delivery due to their ability good biocompatibility, low toxicity, and ability to encapsulate both hydrophilic and hydrophobic drugs. ^1^ Liposomes can also be modified to release drug upon stimuli, such as light or pH. The ability to control the release of a drug provides a pharmacokinetic advantage over passive release systems allowing for tuning dosage as needed for keeping drug concentrations in a therapeutic window. ^2^ Photo-responsive liposomes often utilize a photothermal agent to cause structural changes in the lipid bilayer allowing for drug encapsulated inside the liposome to be released. ^3-8^

This modification can be metal nanoparticles, such as gold nanorods or iron oxide nanoparticles, or organic dyes, such as indocyanine green (ICG). ^9-11^ ICG is a near-infrared (NIR) absorbing organic dye that produces heat when exposed to NIR light that is approved by the FDA. ICG is prone to photobleaching under irradiation making repeated releases difficult. ^3, 12^. Gold nanorods can also undergo structural changes under repeated exposure, reducing their photothermal ability at NIR.^13^ As such the need to develop photostable materials for repeated irradiation exposure is critical to long-term controlled drug release. Croconium dyes are organic dyes formed from a condensation reaction with croconic acid with two molar equivalent electron donating systems.^14^ Croconium dyes poses high NIR absorption, low fluorescence quantum yields, high chemical, thermal, and photo stability, low reactive oxygen species (ROS) production, and synthetic flexibility making excellent potential alternatives.^15^ Croconium dyes provide similar heating levels to gold nanorods at similar levels of absorbance while providing better photostability than ICG which undergoes rapid photobleaching and ROS production which can damage the drug and liposomes making it ideal for photothermal systems.^16^ Croconium dyes have been used in nanoparticle systems for producing a photothermal effect but are uncommon in drug release systems. ^17-21^

Liposomes with sizes in the micron range were synthesized via the reverse-phase evaporation method to allow for an increase in encapsulation efficiency over liposomes produced via lipid film hydration^22^. Our group has previously used this synthesis method for the production of liposomes for integration into a secondary poly(lactide-co-glycolic) acid (PLGA) capsule for a long-term drug delivery system.^2, 23, 24^

In this study we developed a stearic lipid-conjugated croconium dye allowing incorporation into the lipid bilayer for controlled delivery of dexamethasone sodium phosphate (DSP). The study showed effect of laser power and irradiation time on drug release as well as incorporating PCDA into the lipid bilayer for measuring the local lipid bilayer temperature during the photohermal activation upon laser irradiation. This method provided precise local bilayer temperatures over bulk temperature and insight into temperature requirements for drug release. The liposomes displayed a controlled release in the presence of a tissue phantom and showed low cytotoxicity, indicating their potential use in biological applications.

## 2 MATERIALS AND METHODS

### 2.1 Materials

Thiophene-2-thiol, methyl isonipecotate, toluene, croconic acid, chloroform, dichloromethane (DCM), 10,12-pentacosadiynoic acid (PCDA) were purchased from Sigma-Aldrich (St. Louis, MO). Acetic acid, dimethyl sulfoxide, methanol, 1 x phosphate buffered saline were purchased from Fisher Scientific (USA). 1-(3-Dimethylaminopropyl)-3-ethylcarbodiimide hydrochloride (EDC), sulfo-N-hydroxysulfosuccinimide (Sulfo-NHS), MTT (3-(4,5-dimethylthiazol-2-yl)-2,5-diphenyltetrazolium bromide) were purchased from ThermoFisher (Waltham, MA). 1,2-distearoyl-sn-glycero-3-phosphocholine (DSPC), 1,2-distearoyl-sn-glycero-3-phosphoethanolamine-N-[methoxy(polyethylene glycol)-5000] (ammonium salt) (DSPE-Peg5k), 1,2-distearoyl-sn-glycero-3-phosphoethanolamine-N-[methoxy(polyethylene glycol)-2000] (ammonium salt) (DSPE-Peg2k) were purchased from Avanti Reseach (Alabaster, AL). Stearyl amine was purchased from the Tokyo Chemical Industry CO., LTD (Tokyo, Japan). Cholesterol was purchased from MP Biomedicals, LLC. (Solon, OH). Dexamethasone sodium phosphate (DSP) powder was purchased from PCCA (Houston, TX). Human microvascular endothelial cell line-1 was acquired from ATCC (Manassas, VA).

### 2.2 Croconium Dye Synthesis

The synthesis method used in this study was from a report by Song and Foley.^25^. Briefly, an oil bath was heated to 120 °C and 2.32 grams (20 mmol) of thiophene-2-thiol, 4.3 g (30 mmol) of methyl isonipecotate and 20 ml of toluene were placed in a 100 ml round bottom flask. The flask was flushed with nitrogen for 5 minutes. A magnetic stirrer was added to the flask and connected to a water condenser. The flask was placed in the oil bath and refluxed for 2 hours. After 2 hours the reaction mixture was purified with a silica gel column. The effluent was collected and evaporated to collect the desired product, P1. NMR was performed to confirm the desired product. 0.45 g (2 mmol) sample of P1 was added to 10 ml of 0.5 M NaOH was heated to 110 °C and refluxed for 1 hour. The solution was then cooled to room temperature and collected. 10 % acetic acid was added to the refluxed solution to cause a precipitate to form. The product was filtered via gravity filtration and then dried overnight. After drying the desired product, P2, was collected and confirmed via NMR. 422 mg (2 mmol) of P2 and 142 mg (1 mmol) of croconic acid was added to a round bottom flask. 30 ml of a 1:1 volumetric ration of toluene and n-butanol were added to the flask and the solution refluxed for 1 hour at 120 °C. The solution was then cooled to room temperature and the desired product, P3, isolated via gravity filtration. The product was then washed with methanol and dried overnight. The desired product was confirmed via NMR.

### 2.3 Croconium Dye – Steary lamine Conjugation

80 mg of P3 (0.15 mmol), henceforth known as croconium dye (CD) was dissolved in 4 ml of dimethyl sulfoxide (DMSO). 93.14 mg (0.6 mmol) of 1-ethyl-3-(3-dimethylaminopropyl) carbodiimide (EDC) was added to the DMSO solution. The solution was stirred for 3 hours. In a separate vial 40.76 mg (0.15 mmol) of stearyl amine (SA) was dissolved in dichloromethane (DCM). The DCM solution was added to the DMSO solution and 130.28 mg (0.6 mmol) of sulfo-N-hydroxysulfosuccinimide (Sulfo-NHS) added to the solution. The solution was then stirred for 24 hours. The DCM was then evaporated overnight. The remaining solution is added to a 50 ml centrifuge tube and roughly 40 ml of DI water added to the centrifuge tube. The solution was then centrifuged at 10,000 rpm for 10 minutes. The supernatant was discarded leaving the precipitated product. The cycle was repeated until the supernatant is clear. The product was frozen in a -80 °C freezer then dried via lyophilization (Labconco Freeze Dry 4.5). The conjugation of the CD and SA was confirmed via FTIR and UV-Vis.

### 2.4 Liposome Synthesis

A mixture of 1,2-distearoyl-sn-glycero-3-phosphocholine (DSPC), 1,2-distearoyl-sn-glycero-3-phosphoethanolamine-N-[(polyethylene glycol)-5000] (DSPE-PEG5k), cholesterol, and stearyl amine (SA), with a mole ratio of 50:5:35:10, dissolved in chloroform was added to a round bottom flask. A rotary evaporator was used to evaporate the chloroform to form a thin film of lipids in the flask. The film was then rehydrated using 1.5 ml of a 2:1 volumetric ratio of chloroform:methanol. For synthesis of CD-SA liposomes, SA was substituted with CD-SA. Dexamethasone sodium phosphate (DSP) (103.25 mg, 200 μmol) in 1.5 ml of 1 x PBS was added dropwise to the lipid mixture in an ice bath while stirring. The aqueous - organic mixture was stirred for 30 minutes. The flask was then removed from the ice bath and sonicated for 10 minutes. The organic phase was removed using a rotary evaporator placing the solution under a vacuum for 30 minutes. The remaining phase was collected and vortexed for 20 minutes. The liposomes were then purified using a PD-10 column followed by a 72 hours dialysis changing the buffer every 8 hours. Finally, the liposomes were centrifuged at 5,000G for 1 hour and the supernatant was replaced with 1 x PBS to a total final volume of 3 ml. The liposomes were then characterized via transmission electron microscopy (TEM), dynamic light scattering (DLS), and UV-Vis spectroscopy. The details are provided in our previous publication. ^2^

### 2.5 Variable Power Drug Release

The liposomes (100 μL) from section 2.4 were placed under a 808 nm laser (LSR808NL-2W) at currents of 440, 510, and 580 mA. The power at each current was recorded using a power meter (843-R, Newport Optics). The resulting power was 210, 295, and 380 mW respectively (4.25, 6.0, 7.75 W/cm^2^). The liposomes were placed under the 808 nm laser for 0, 1 and 2 minutes. The liposomes were then allowed to sit overnight at room temperature. The filtrate was separated from the liposomes using 10 kDa amicon filters. The liposomes were centrifuged at 14,000 G for 20 minutes to separate drug released in the supernatant. The drug concentration was analyzed using UV-Vis spectroscopy with a calibration curve of DSP.

### 2.6 PCDA Liposome Synthesis and Temperature Ladder

10,12-Pentacosadiynoic acid (PCDA) was additionally added to the lipid composition of the liposomes at a mole ratio 25:5:35:25:10 of DPSC, DSPE-PEG2K, cholesterol, PCDA, and SA/CD-SA. The liposomes were synthesized following the method described in section 2.4. The liposomes were subsequently exposed to 30 W 254 nm UV-light for 30 minutes to polymerize the PCDA, resulting in a blue coloration. The liposomes were heated in an oil bath at the desired temperature (35-75 °C) for 1 minute, resulting in different degrees of red shift in UV-Vis spectrum. The liposomes were then characterized via UV-Vis spectroscopy.

Using trapezoidal integration and the equations 1-3, the degree of this red shift was quantified.

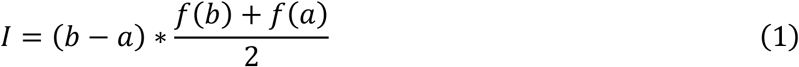

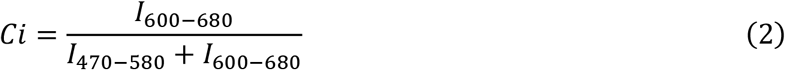

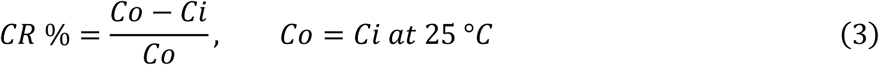

Plotting the values a sigmoidal function emerges

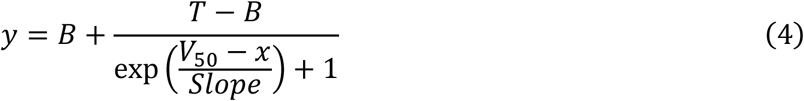

where B is the bottom asymptote, T is the top asymptote, V_50_ is the inflection point, and slope describes the steepness of the curve. Polymerized liposomes were irradiated at the powers and times as described in section 2.5. Th UV-Vis spectrum of the irritated liposomes were analyzed using the same trapezoidal integration method to determine the CR %. Using the sigmoidal function the local liposome temperature was then estimated.

### 2.7 Agar Phantom Tissue Drug Release

Simulated tissue phantoms were made to determine the ability of the liposomes to release drug beneath tissue^26^. The tissue phantoms were made by sodium alginate, 0.1 v% of India ink, and 1 wt% or 3 wt% agar was added to the solution. The solution was then heated to a boil then allowed to cool slightly. The warm solution was poured into 3D printed molds of various depths, 1, 3, 6 mm. Mold was then covered with a slide to press the solution to the exact height. The molds were allowed to rest until the gel solidified. 100 μl of liposomes prepared the using the same methods in section 2.4 were placed under the 808 nm laser at 380 mW with phantom tissue thickness of 0, 1, and 3 mm placed over top the liposomes. The liposomes were irradiated for 2 and 5 minutes. The liposomes were then allowed to sit overnight at room temperature. The filtrate was separated from the liposomes using 10 kDa amicon filters. The liposomes were centrifuged at 14,000 G for 20 minutes. 50 μl of filtrate was collected for UV-Vis analysis.

### 2.8 MTT Cytotoxicity

HMEC-1 cells were used to test the cytotoxicity of CD-SA liposomes. The cells were seeded in a 96 well plate at 10,000 cells per well in 100 μl of growth medium. The cells were incubated for 24 hours at 37 °C in 5 % CO_2_. After 24 hours the growth medium was removed from the cells and replaced with starvation medium and starvation media mixed with the liposomes at various croconium dye concentrations. The croconium dye concentrations in the media was 250, 100, 50, and 25 μM. The cells were incubated for 24 h at 37 °C in 5% CO_2_. The starvation media/ starvation media with liposomes was removed from the cells and MTT dissolved in starvation was added to the cell. The MTT starvation media was allowed to incubate for 3 hours at 37 °C in 5% CO_2_. The MTT solution was removed and 100 μl of DMSO was added to the well and mixed well. The well plate was placed back in the incubator for 10 minutes. The absorbance at 550 nm was then recorded. The thermal effect of the croconium dye was also tested. Cells were incubated for 24 hours with growth media in a 96 well plate. Starvation media with a croconium dye concentration of 250 μM was added to the well. The cell were then irradiated with an 808 nm laser at 380 mW for 1, 2, and 5 minutes. After irradiation the cells were incubated for 30 minutes to stabilize in the 37 °C in 5% CO_2_ environment. The starvation media was removed and replaced with MTT starvation media an incubated for 3 hours. The MTT solution was removed and 100 μl of DMSO was added to the well and mixed well. The well plate was placed back in the incubator for 10 minutes. The absorbance at 550 nm was then recorded.

## 3 RESULTS

### 3.1 Croconium Dye – Stearyl Amine (CD-SA) Conjugation

The conjugation of croconium dye to stearyl amine was confirmed by FTIR. Peaks at ∼ 2915 and 2850 cm^-1^ present in stearyl amine along with peaks at ∼ 1680 cm^-1^ in croconium dye were confirmed in CD-SA (**Figure 2A**). The UV-Vis spectrum also showed that the product in chloroform has a characteristic peak of CD at 780 nm (**Figure 2B**). This also indicated successful lipid conjugation as the unconjugated dye is insoluble in chloroform.

**Figure 1:**
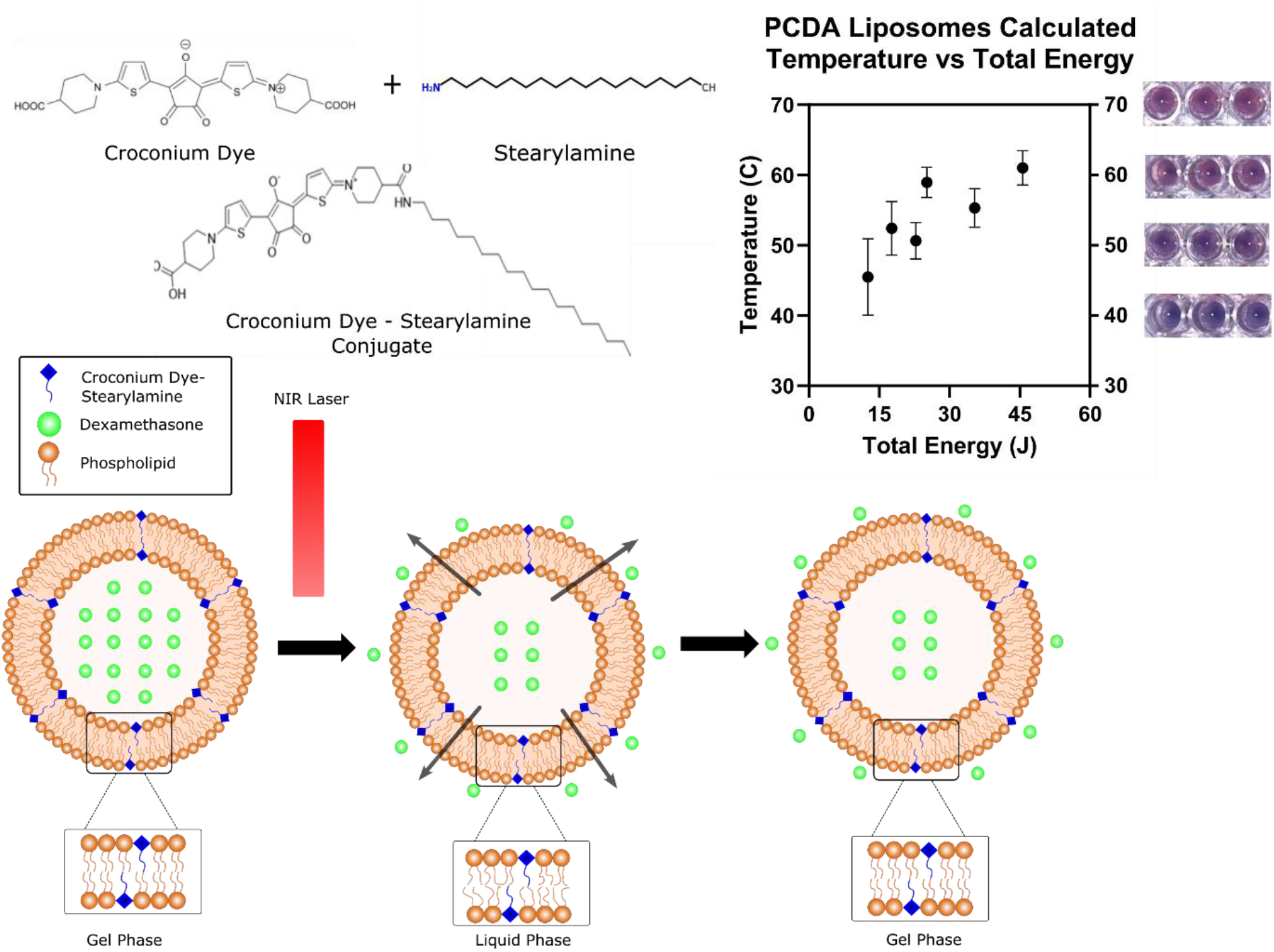
Overall Schematic

**Figure 2A.**
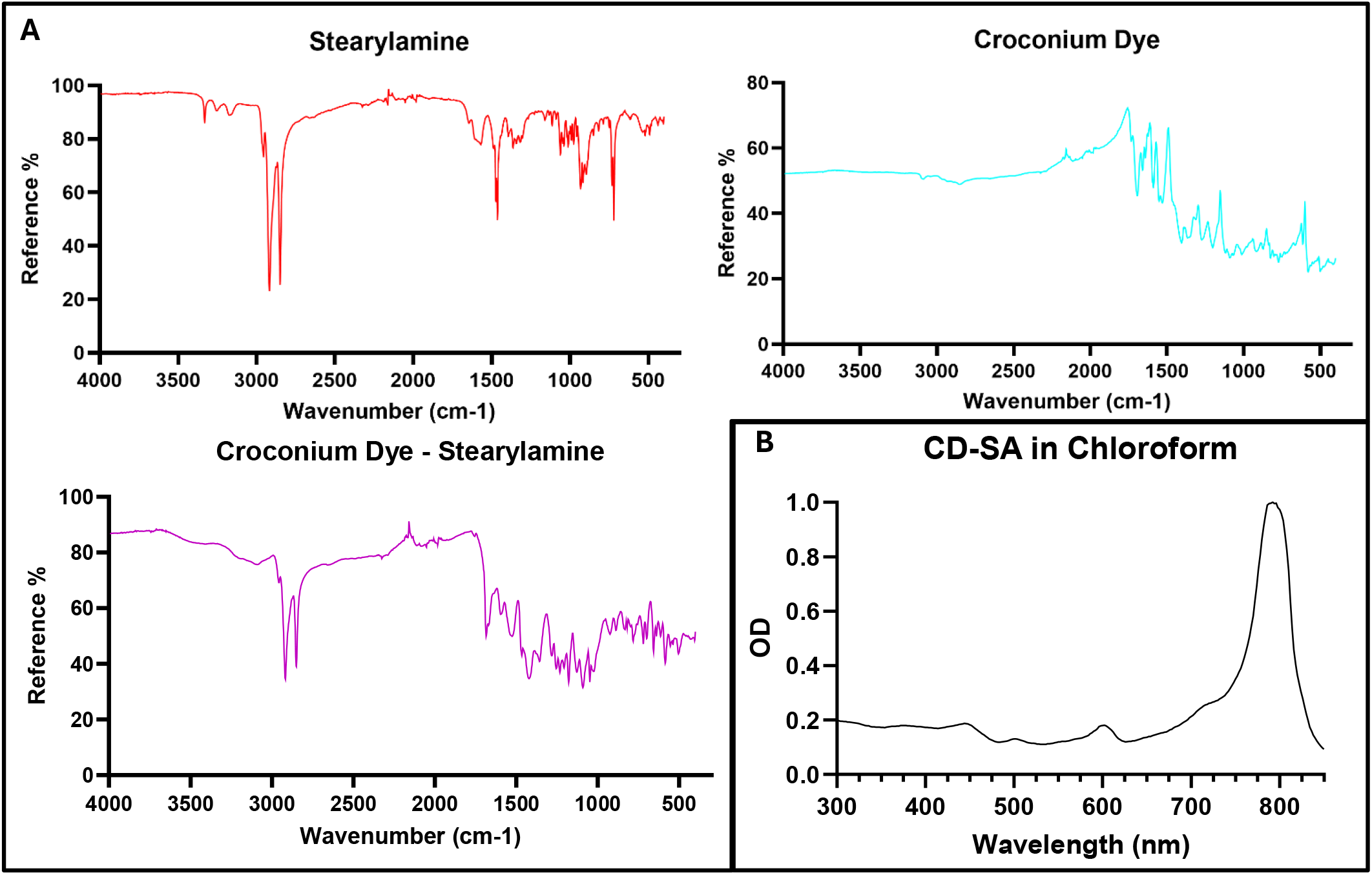
FTIR Spectrums Figure 2B. CD-SA UV-Vis Spectrum (Chloroform)

### 3.2 Characterization of SA and CD-SA Liposomes

The liposomes had an effective diameter of 700 nm for the stearyl amine (SA) liposomes and 630 nm for the croconium-dye stearyl amine (CD-SA) liposomes, based on the DLS intensity average (**Figure 2A**). This was supported by TEM imaging in size of ∼ 600 nm for the CD-SA liposomes shown in **Figure 2B**. The CD-SA liposomes showed two peaks one at 650 nm and the other at 780 nm. The 650 nm peak is observed only when CD-SA is embedded within the lipid bilayer. This behavior is likely due to nanoconfinement of the dye within the membrane, resulting in a blue-shifted absorption band. When conjugated to stearic acid, the CD molecules are expected to be closely packed within the bilayer, enabling electronic interactions that split the original transition into multiple absorption bands. The partial retention of the 780 nm peak suggests that a fraction of the dye remains exposed to the aqueous environment, while the rest resides in a more ordered and constrained membrane phase. The lipid bilayer restricts molecular motion and conformational flexibility, favoring specific orientations that lead to these spectral changes.^27 28^

### 3.3 808 nm Laser Drug Release

The CD-SA liposomes successfully released DSP when irradiated by the 808 nm laser whereas the SA liposomes showed no release under irradiation (**Figure 4**). Upon irradiation at 210 mW the CD-SA liposomes released 0.92 μg and 4.83 μg after 1 and 2 minutes, respectively. At 295 mW the released amounts increased to 3.19 μg and 10.19 μg for 1 and 2 minutes, respectively. At 380 mW the liposomes released 7.13 μg and 18.84 μg after 1 and 2 minutes, respectively. One-way Anova was performed to test the statistical significance of the release. For the 210 mW irradiation the p-values were 0.4276 for 1-minute and < 0.0001 for the 2-minute, compared to 0 min. The results indicate that 2-minute irradiation at 210 mW resulted in a statistically significant level of drug release. For the 295 mW irradiation the p-values were 0.0304 for 1-minute and < 0.0001 for the 2-minute compared to 0 min and for the 380 mW irradiation the p-values were 0.0110 for the 1-minute and < 0.0001 for the 2-minute compared to 0 min, indicating that the release was also power-dependent in statistical significance of drug release. Based on the release amounts, 4.83 μg released after 2 min at 210 mW could be clinically relevant, especially for DSP ocular local delivery. The bulk temperature change of the liposomes was recorded using a photothermal camera. The bulk temperature of the SA liposomes showed no change in temperature. The 1-minute CD-SA liposomes showed a change of 3.7, 4.7, and 6.0 °C for the 210, 295, and 380 mW, respectively. For the 2-minute irradiation the CD-SA liposomes showed a change of 5.9, 8.8, and 11.9 °C, respectively. The bulk temperature changes for all irradatioin times and power resulted in a p-values < 0.0001. The bulk temperature heating curve can be seen in the supplementary information (**SF2**)

**Figure 3A.**
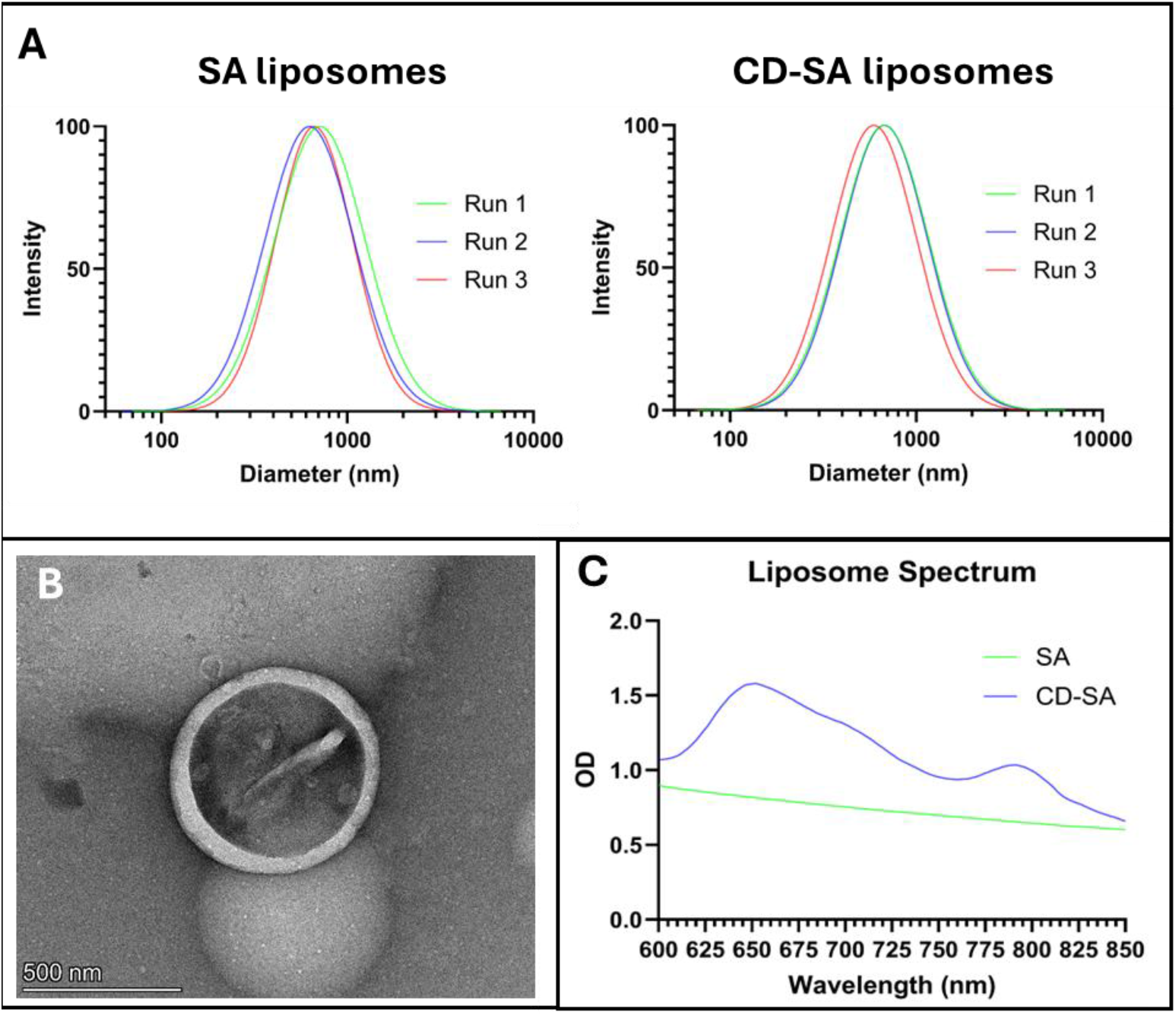
DLS intensity size distribution of SA liposomes and CD-SA liposomes; B. TEM of CD-SA liposomes; C. UV-Vis spectra of SA liposomes and CD-SA liposomes.

**Figure 4.**
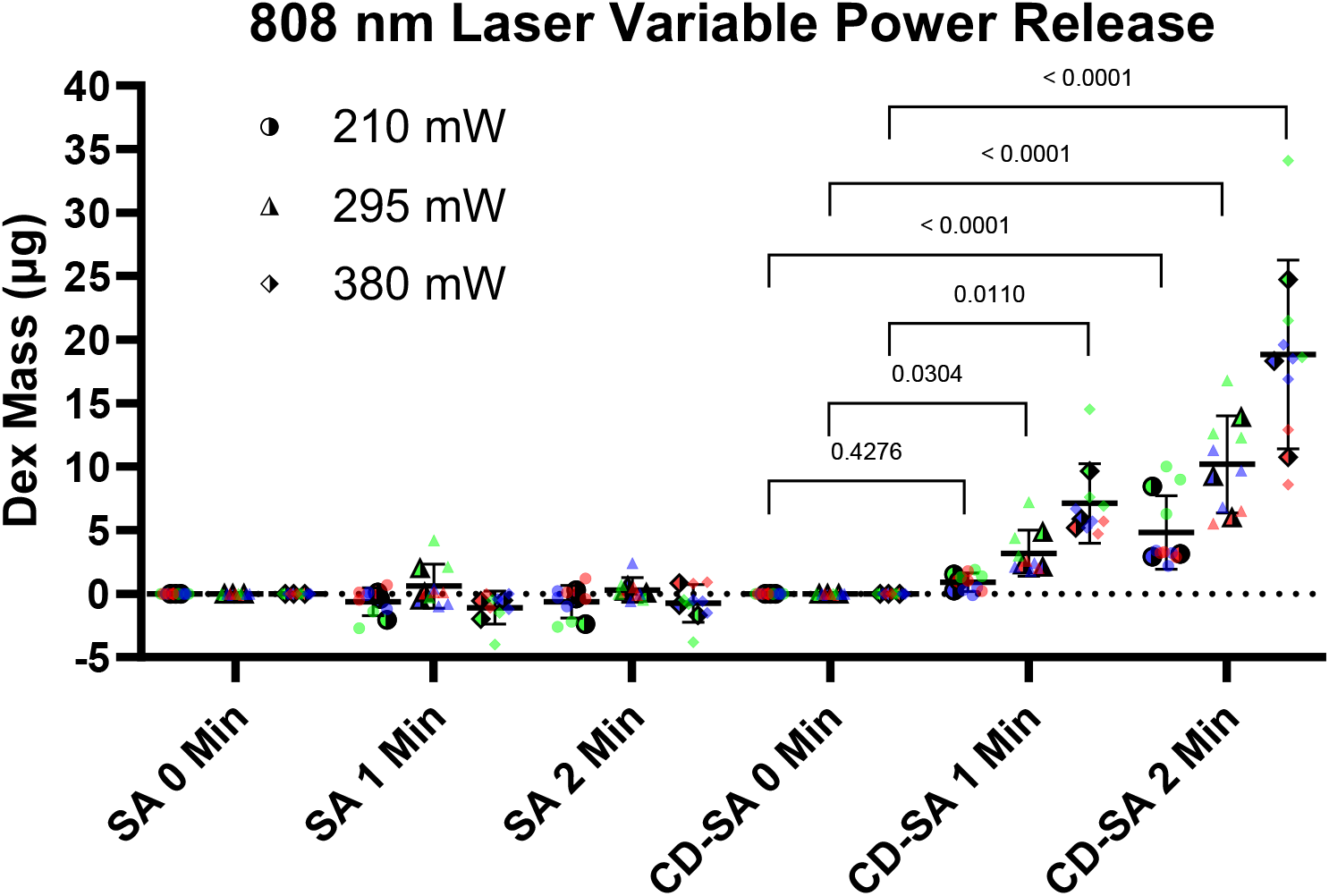
Drug release from CD-SA liposomes compared to SA liposomes at various powers and durations of laser.

### 3.4 PCDA Liposome Local Average Temperature

From the bulk temperature changes from section 3.3 it was shown that a ΔT of 4.7 °C was the minimum temperature to achieve statistically significant drug release. This was an average final bulk temperature of 31.2 °C. However, the known liquid phase of DSPC is 55 °C^29^. As such there was a desire to more accurately determine the local temperature of the liposome bilayer under irradiation to better understand the liquid phase transition in the liposomes for the drug release.

This was done through the incorporation of PCDA into the lipid bilayer of CD-SA liposomes and SA liposomes. PCDA has been broadly used a temperature sensor due it its ability to perform a colorimetric shift from blue to red when heated^30^. To determine the local bilayer temperature a temperature calibration curve of the polymerized PCDA liposomes was established by heating the liposomes at various temperatures. The liposomes showed various degrees of red shift in the spectrum (**Figure 5A,B**). The greater the temperature of the oil bath the greater the degree of red shift. Due to the croconium dye in the CD-SA liposomes the absorbance in the 650 nm region was increased over the SA liposomes. The CD-SA liposomes also showed a greater degree of polymerization likely due to better alignment and packing the CD-SA liposomes due to the addition of croconium dye in the bilayer. The quantitative shift was determined using CR% values (please see section 2.6 for CR% calculation).

**Figure 5.**
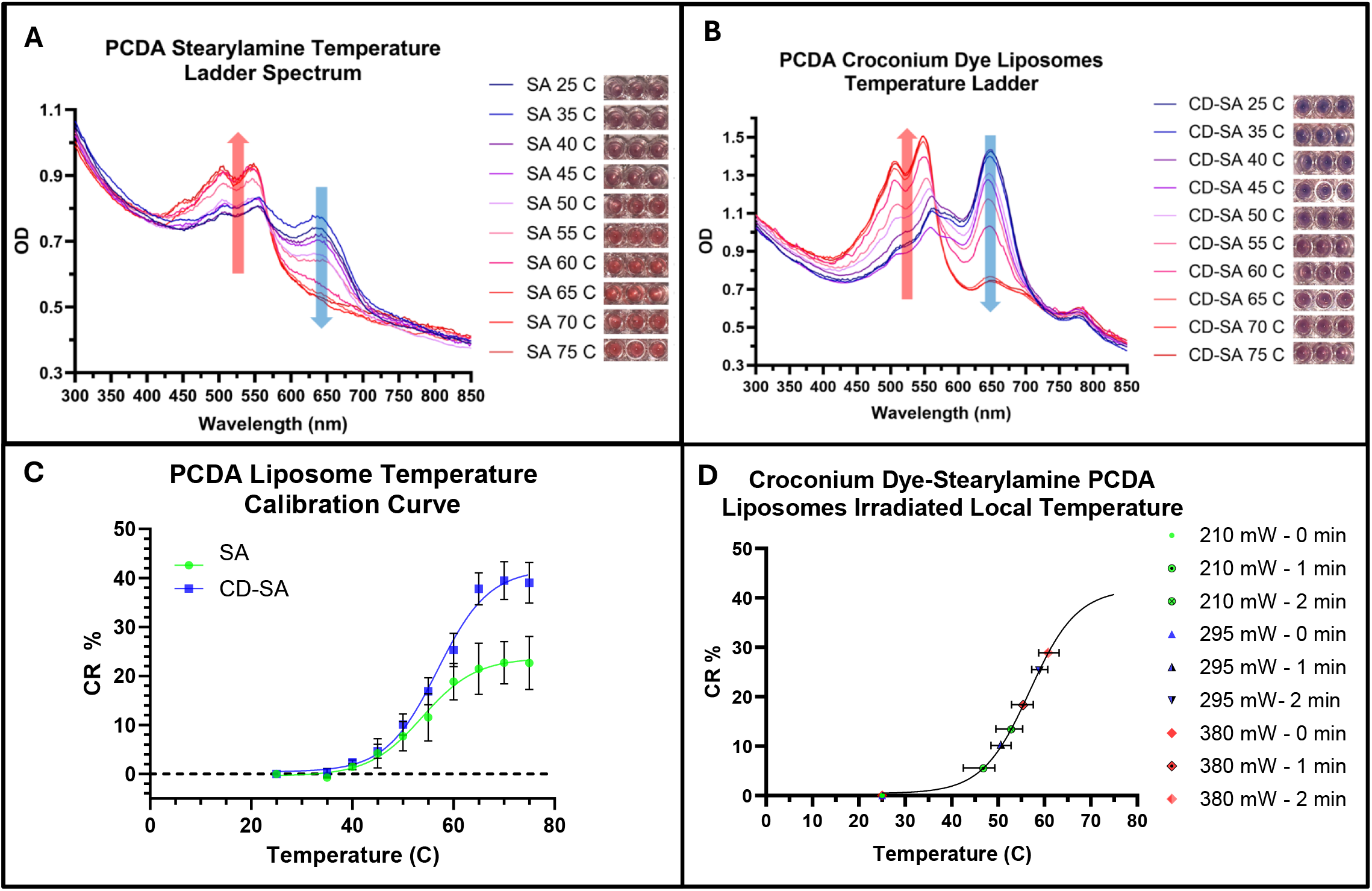
UV-Vis spectra of heated PCDA liposomes at various temperatures for A. SA liposomes, B. CD-SA liposomes, C. Temperature calibration curves for SA liposomes, and CD-SA liposomes, D Local temperature of lipid bilayer upon laser irradiation, estimated using the CR % curve.

The average CR % values for heated temperatures of 35 – 75 °C ranged from -0.7 to 22.7 and 0.3 to 39.0 % for the SA and CD-SA liposomes respectively (**Figure 5C**). Plotting the values a sigmoidal function emerges, 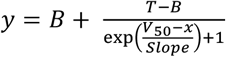. The values from each of the parameters of the regression models can be seen in Table 1. The regression model for the PCDA-SA liposomes had an R^2^ value of 0.76 while the regression model for the PCDA-CD-SA liposomes had an R^2^ value of 0.94. The PCDA-SA model had greater variation in its polymerization between trials which resulted in a lower R^2^ value though all trials followed a sigmoidal regression.

**Table 1:** Parameters for Sigmodial Regression Model for PCDA Temperature Calibration.

| Parameter | Stearyl amine Liposomes | Croconium dye – stearyl amine Liposomes |
| --- | --- | --- |
| B | -0.39 | 0.43 |
| T | 23.67 | 41.82 |
| $V_{50}$ | 53.83 | 56.81 |
| Slope | 5.29 | 5.10 |

Polymerized liposomes were irradiated at the current and times as described in section 3.3. The average CR % were 5.5, 13.4, 10.3, 25.3, 18.3, 28.9 % for the 210 mW, 1- and 2-minute irradiation, 295 mW 1- and 2-minute irradiation, and the 380 mW 1- and 2-minutes irradiation respectively. Using the equation established above the average local temperature of the liposomes were calculated as 46.8, 52.8, 50.6, 58.9, 55.4, 60.8 °C respectively (**Figure 5D)**. This showed for our liposomes system to achieve a statistically significant drug release the liposomes needed reach a bilayer temperature of 50 °C a temperature slightly less than the known liquid transition temperature for DSPC of 55 °C.

### 3.5 808 nm Laser Drug Release with Tissue Phantom

The CD-SA liposomes were irradiated in the presence of tissue phantoms to account for attenuation of light due to scattering and absorption at varying tissue types and depths. Agar gels of 1% and 3% with 0.1% India ink were used to mimic adipose/connective tissues (μ = 0.1 – 0.7 mm^-1^) and dermis tissue (μ = 0.5 – 1.5 mm^-1^), respectively. ^26, 31-35^ The attenuation coefficients of the 1% agar and 3% agar tissue phantoms were μ= 0.493 mm^-1^ and 0.651 mm^-1^. The 380 mW power was used for all phantom tissues. For the non-tissue control the average release was 19.1 μg and 37.5 μg for the 2-minute and 5-minute irradiation, respectively (**Figure 6**). The 2 minute data was consistent with the data from **Figure 4**. For the 1 mm tissue thickness with 1 % agar composition the average mass release was 2.2 μg and 4.9 μg for the 2-minute and 5-minute, respectively. For the 1 mm tissue thickness with 3 % agar 1.1 μg and 2.7 μg for the 2-minute and 5-minute irradiation, respectively.

**Figure 6.**
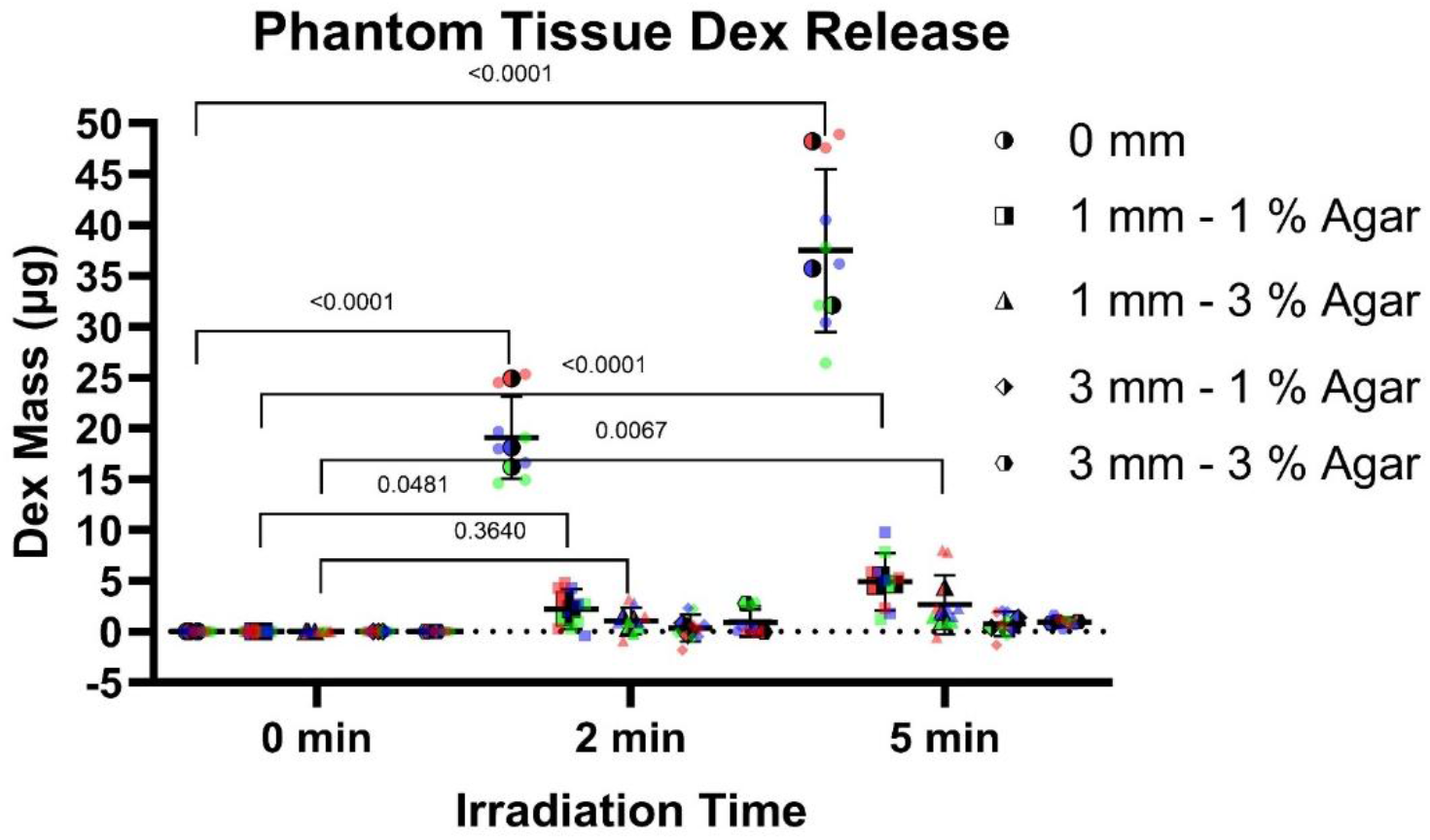
Drug release in the presence of tissue phantoms to account for different tissue types and depths.

For the 0 mm tissue thickness the p-values between 0 min and 2 min, and between 0 min and 5 min were both less than 0.0001, indicating that the effect of laser duration on the drug release was statistically significant when the liposomes were directly exposed to the laser without light attenuation by tissue phantom. For the 1 mm, 1 % agar tissue phantoms showed statistically significant release compared to 0 min for both irradiation times with a p-value of 0.0481 for the 2-minute irradiation and < 0.0001 for the 5-minute irradiation. For the 1 mm, 3 % agar the 2-minute irradiation did not show a statistically significant release having a p value of 0.3640 but did show a statistically significant release for the 5-minute irradiation having a p value of 0.0067. The results suggest that longer laser irradiation may be needed to activate drug release when the light attenuation is significant depending on the tissue type. For the 3 mm tissue thickness there was no statically significant release for either the 1 or 3 % agar.

The bulk temperature change of the liposomes, for the no tissue phantom, was 9.8 and 15.7 °C for the 2-minute and 5-minute, respectively. For the 1mm, 1 % agar the bulk temperature change was 4.5 and 5.8 °C for the 2-minute and 5-minute respectively. The 1 mm, 3 % agar showed reduced bulk heating at 3.3 and 4.7 °C for the 2-minute and 5-minute respectively. The 3 mm tissue showed an increase of 1.9 and 3.3 °C for the 1 % agar and 1.3 and 2.3 C for the 3 % agar for the 2- and 5-minute irradiation times respectively (**SF3**). All bulk heating showed a statistically significant p-value with all p-values being < 0.0001.

Based on the results potentially clinically significant release is possible but optimization is likely necessary to improve absorbance for release in the presence of tissue. One such improvement could be the use of a pulsed laser as they show increased tissue penetration in comparison to continuous wave lasers while reducing tissue heating.^36^

### 3.6. Safety against cytotoxicity and the bulk heating

The cytotoxicity of the CD-SA liposomes was determined based on MTT absorbance, exploring the chemical and thermal cytotoxicity of the liposomes (**Figure 7A,B)**. Chemical cytotoxicity was evaluated at four CD-SA liposome concentrations (250, 100, 50, and 25 μM) using cells without liposomes as the control. CD-SA liposomes did not negatively affect cell viability; in fact, increasing CD-SA liposome concentration corresponded to higher viability, with values of 162%, 127%, 115%, and 110% at 250, 100, 50, and 25 μM, respectively. The CD-SA liposomes showed a statistically significant increase in cell growth for the 250 μM and 100 μM concentration with p-values of <0.0001 and 0.0259. There was no statical difference in the 50 and 25 μM concentration with p-values of 0.2659 and 0.5476

**Figure 7A.**
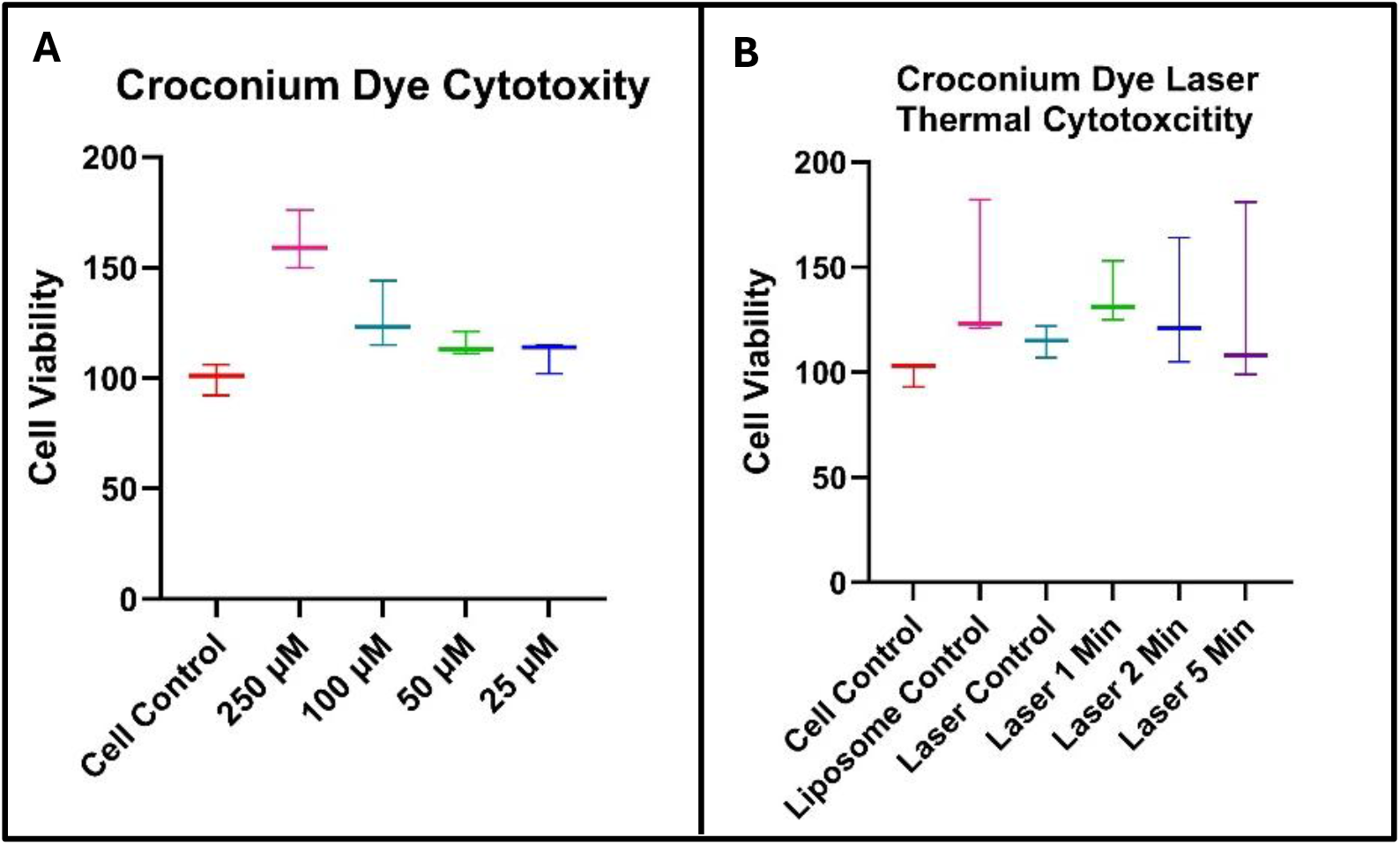
Cytotoxity of CD-SA liposomes, 7B. Thermal Toxicity of CD-SA liposomes under irradation

Thermal cytotoxicity was assessed by irradiating CD-SA liposomes at 250 μM at 380 mW for 1, 2, and 5 minutes. Cells without liposomes were also irradiated for 5 minutes as a laser control. The liposome-only control exhibited 142% viability, while the laser-only control showed 115% viability. Irradiated liposome samples maintained high viability, with values of 136%, 130%, and 129% following 1-, 2-, and 5-minute irradiation, respectively. There was no staticial difference in the cell control and any other group with p-values of 0.2622, 0.9365, 0.3779, 0.5420, and 0.5609

In summary, no chemical and thermal toxicity of the CD-SA liposomes was observed with and without laser irradiation.

## 4 CONCLUSION

Our study comprehensively examined release behavior and cytotoxicity of a novel liposome drug delivery system through *in-vitro* assessments. The liposomes showed stimuli responsive drug release and the linking of liposome bilayer local temperature to drug release provides an increased understanding of the liposome bilayer transition from a gel to liquid phase and its effect on drug release. The findings show the effectiveness of the liposomes as an on-demand drug delivery platform with potential for integration for long term drug delivery.

## 5 SUPPLEMENTARY

Bulk Temperature Heating Graphs for Variable Power and Phantom Tissue

