## Supplemental Information for "Croconium Dye Thermosensitive Liposomes for Light Activated Drug Release"

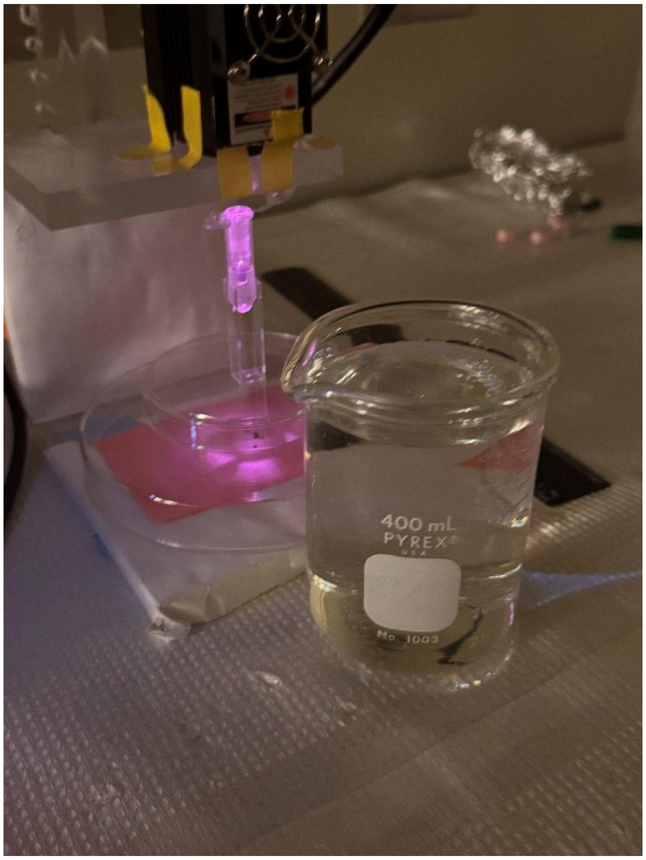


**SF1.** 808 nm Laser Setup

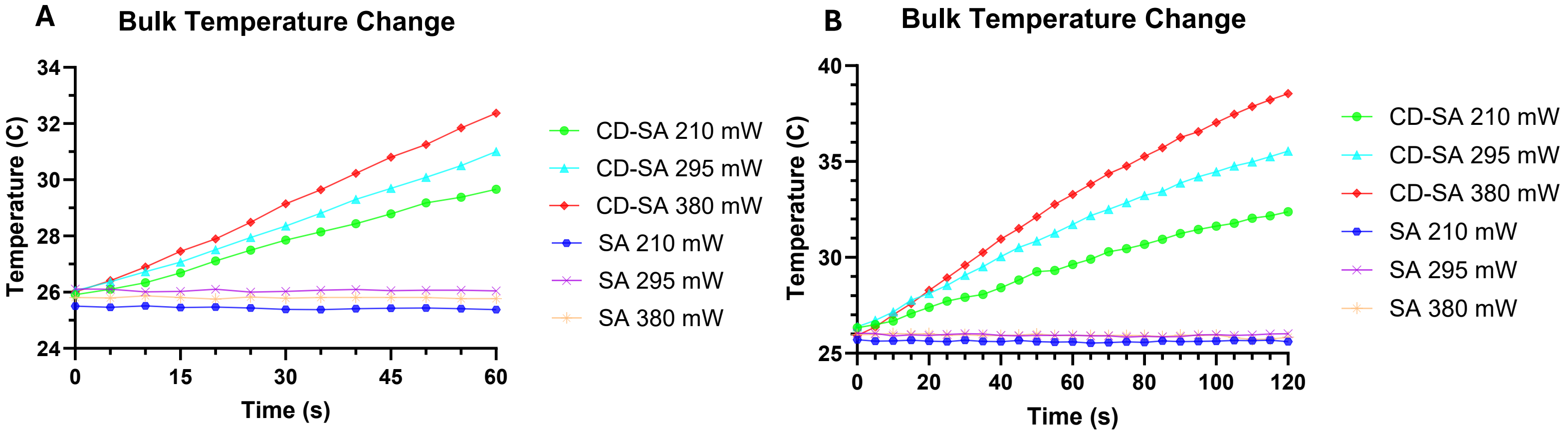


**SF2A.** 1 Minute Bulk Temperature **B.** 2 Minute Bulking Temperature

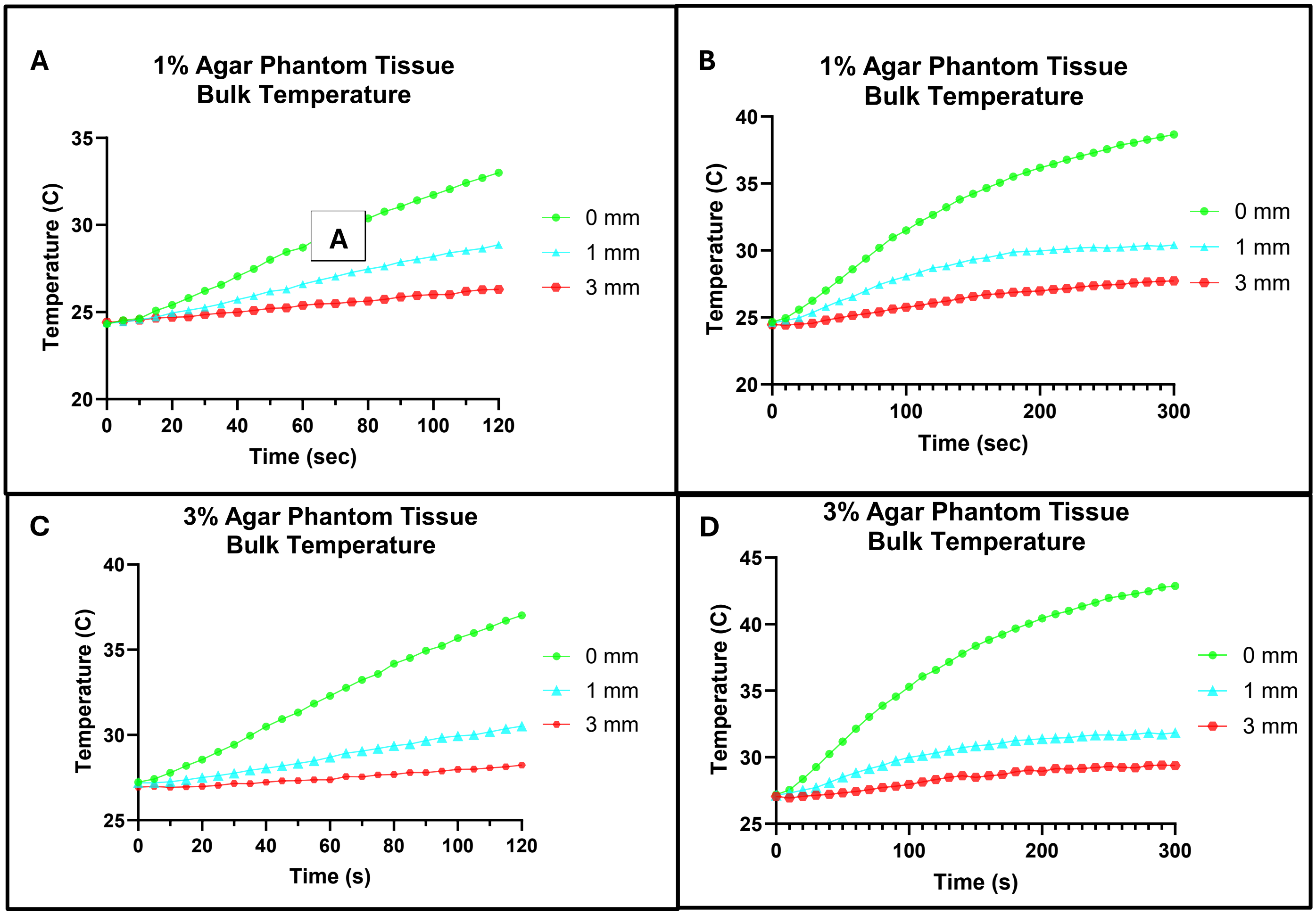


**S3A.** 1 % 2-Minute Bulk Temperature **B.** 1 % 5-Minute Bulking Temperature
**C.** 3 % 2-Minute Bulk Temperature **D.** 3 % 5-Minute Bulking Temperature
